# Intracellular auto-circularizing RNA using human tRNA splicing elements

**DOI:** 10.1101/2025.07.28.667085

**Authors:** Qianbo Ma, Chuanqing Tang, Jie Wang, Yingchun Liu, Xiulian Sun, Sheng Zhao

**Affiliations:** Department of Neurology, Shandong Key Laboratory of Mitochondrial Medicine and Rare Diseases, Research Institute of Neuromuscular and Neurodegenerative Diseases, Qilu Hospital of Shandong University, Jinan, Shandong, China; HomiX Biotechnology Co., Ltd. Nanjing, China; uBriGene Biosciences Co., Ltd. Beijing, China

**Keywords:** Circular RNA, human tRNA splicing elements, auto-circularizing RNA

## Abstract

Circular RNAs (circRNAs), a type of covalently closed RNA, have emerged as crucial regulatory factors in gene expression and development across various organisms. Understanding their formation and functional mechanisms holds potential for therapeutic applications, including novel RNA vaccines against diseases such as COVID-19. However, practical challenges still exist in the preparation of circRNAs. In this study, we explored the influence of human tRNA splicing elements on circRNAs formation and identified effective sequences for intracellular auto-circularization. We developed a generic auto-circularizing RNA (acRNA) framework template, optimized various key components of circRNAs formation, and proposed a more efficient and precise intracellular auto-circularization method. This study provides novel insights into the production mechanism of circRNA and establishes a simple and effective method for intracellular auto-circularization.

## Introduction

Circular RNAs (circRNAs) are a captivating class of RNA that exert significant influence over gene expression and developmental processes across diverse organisms^[1,2]^. These unique molecules are formed through a back-splicing event, which fuses the 3′ and 5′ ends of RNA to form closed-loop structures^[3,4]^. They have been identified across the evolutionary spectrum, spanning from archaea to human, and play pivotal roles in numerous biological processes^[5]^. Recent investigations have demonstrated circRNAs could regulate gene expression and protein synthesis through different mechanisms^[2,6]^. For instance, they can act as miRNA sponges to specifically inhibit the binding of certain miRNAs to their target mRNAs; or modulate gene transcription levels by interacting with transcription factors^[7-9]^. Moreover, owing to the remarkable stability and low immunogenicity, circRNAs have emerged as a highly coveted target in the field of novel RNA vaccines^[10]^. Especially during the current COVID-19 outbreak, more and more researchers have begun to develop effective prevention and treatment strategies against infectious diseases using circRNA^[11]^.

Currently, the synthesis of circRNA primarily includes chemical ligation, enzymatic ligation, and ribozyme-assisted synthesis with specially designed sequences^[12-14]^. Among these methods, permuted intron-exon (PIE) is one of the most extensively researched and utilized techniques for synthesizing circRNA, especially for large circRNAs which were used for protein expression^[15]^. Although PIE has been widely used and confirmed to produce exogenous circRNA for stably and efficiently protein expression in eukaryotic cells, it suffers from drawbacks such as low circularization efficiency and by-products^[16-18]^, resulting expensive price for manufacture and challenge for quality control. Recent studies have successfully synthesized small circular RNA aptamers *in vitro* or in cells by utilizing the Metazoans tRNA intron, the Tornado (Twister-optimized RNA for durable overexpression) system^[19]^. However, this method is currently limited to short RNA aptamers such as Broccoli, which are only 49 nt in length. No reports have been conducted to use Tornado system to circularize the long RNA for protein expression.

Additionally, Tornado system employs partial non-human sequences from Metazoan tRNAs, particularly the intronic region, which might raise the concern of potential safety risks for future clinic application.

Here we screened the sequences at splicing site (exonic or intronic) in human tRNAs with introns for their abilities to forming large circRNA intracellularly. The effective sequences from these human tRNA were proved to function as auto-circularizing elements to produce circRNA automatically using the endogenous tRNA splicing machinery. This study provides a novel tool kit of engineered auto-circularizing RNA (acRNA) to generate cellular circRNA for gene expression, which is a cheaper and simpler manufacturing technology for circRNA.

## Materials and Methods

### Plasmid construction

The plasmid backbone with multiple cloning sites was utilized for subcloning the sequences containing human tRNA splicing elements, resulting in a novel vector cassette harboring coding gene. Specifically, the cassette starts with a T7 promoter followed by a 5′ self-cleaving ribozyme containing human tRNA 3’-exonic or 5’-intronic sequence near tRNA splicing site, an internal ribozyme entry site (IRES), gene of interest (e.g. Neongreen-teLuciferase reporter or CRISPR-Cas9 protien), a 3′ self-cleaving ribozyme containing human tRNA 5’-exonic or 3’-intronic sequence near tRNA splicing site, and a single restriction endonuclease recognition site for plasmid linearization (Fig.1). The sequences of the self-cleaving ribozyme containing human tRNA exonic or intronic sequence near tRNA splicing site were shown in supplemental table 1. The self-cleaving ribozyme were designed to generate the tRNA exonic or intronic ends after self-cleavage on both sides of the auto-circularizing RNA (acRNA).

### acRNA synthesis

The *in vitro* transcription (IVT) template were prapared by linearizing the plasmid using the single restriction endonuclease followed by purification using TIANquick midi kit (TIANGEN, Beijing, China). Next, RNAs were synthesized from the linearized template using Hi-yield T7 IVT reagents (Hzymes Biotech, Wuhan, China) for 3 hours. During the IVT process, the ribozymes automatically cut themselves on both ends to generate the acRNA with 5’-OH and 3’-2,3-cyclic phosphate. The IVT reactions were then incubated with DNase I (Hzymes Biotech, Wuhan, China) for 30 mins to remove the plasmid templates. Finally, acRNAs were purified by lithium chloride (LiCl, Invitrogen) precipitation.

### Cell culture and Transfection

HEK293T, CHO and HeLa cells were purchased from American Type Culture Collection (ATCC). Cells were cultured with DMEM medium containing 10% fetal bovine serum (FBS, Lonsa, Shanghai, China) and 1× penicillin / streptomycin antibiotics (BasalMedia, Shanghai, China). Cells were cultured in an incubator at 37°C with 5% CO_2_ and 90% relative humidity.

RNA delivery was achieved by the Ubri-PEI (Ubrigene, Suzhou, China) transfection. The day before transfection, 2×10^5^ cells per well were seeded in 24-well plates. When the cell confluence reached 70%, the obtained acRNAs containing reporter genes were transfected into HEK293T, CHO or HeLa cells using PEI according to the manufacturer’s instructions. Green fluorescence was observed by fluorescence microscopy at 24h and 48h after transfection, and follow-up tests were performed.

### Luciferase assays

After 48 h RNA transfection by PEI, cells were digested from the 24-well plate using trypsin. The cell concentration was measured and then adjusted to 4×10^5^ cells/ml. Next, cells were transferred to 96-well flat-bottom white plates with 90 μl per well. Each experimental group was provided with three replicate wells. Triton X-100 (10%, 10 μl per well) was added to wells and incubated at room temperature for 5-10 min. Finally, the bioluminescence signal were read by a microplate bioluminescence reader equipped with substrate injectors. The final concentration of bioluminescence substrate diphenylterazine (DTZ) was 30 μM in 50 mM Hepes.

### *In vivo* RNA circularization detection assays

After 48 h RNA transfection by PEI, total RNAs were extracted from cells by using RNA-easy isolation reagent (Vazyme, Nanjing, China). cDNAs were obtained by reverse transcription of RNAs using HiScript II strand cDNA synthesis kit (Vazyme, Nanjing, China) according to manufacturer’s instructions and specific reverse transcription primer for the acRNA was used. PCR amplification was then performed and the amplified fragments were detected using agarose gel electrophoresis, recovered using universal DNA Purification kit, and sequenced to confirm the circularization of the acRNA. Sequences of all primers were listed in supplemental table 2.

### Evaluation of the intracellular immune response caused by RNA

The same amount of acRNA, circRNA (in vitro circularized based on PIE method), or linear mRNA were transfected into HEK293T cells by using PEI. According to the method described above, after transfection for 48h, Total RNAs were extracted from cells by using RNA-easy isolation reagent. cDNAs were obtained by reverse transcription of RNAs using HiScript II strand cDNA synthesis kit according to manufacturer’s instructions and Random hexamers was used. Subsequent RT-qPCR was performed using AceQ qPCR SYBR Green Master Mix (Vazyme, Nanjing, China). Data were collected with the CFX96 instrument (BioRad). After obtaining RIGI, CCL2, TNF, EIF2AK2 and GAPDH sequences from NCBI website, RT-qPCR primers were designed using Primer premier 6.0 software. All RT-qPCR primers were synthesized by General Biology (Chuzhou, China), and primers sequences were shown in supplemental table 2. GAPDH was the control, and the relative mRNA expression was calculated by 2^ΔΔ Ct^.

### Detection of *in vivo* gene-editing activity of acRNA containing Cas9 protein

To investigate whether long sequence (>5 kb) acRNAs can be successfully circularized in vivo, we constructed plasmids containing Cas9 protein. To verify the activity of Cas9 protein, we constructed a HEK293 cell oveexpression Neongreen fluorescent protein fused te-luciferase. Successful gene editing using sgRNA targeting the Neongreen coding sequence would result the reduction of fluorescence and bioilluminance signal. RNA delivery was achieved by lipofectamine transfection. The day before transfection, 2×10^5^ cells per well were seeded in 24-well plates. When the cell confluence reached 70%, Cas9-acRNA and sgRNA agaist NeonGreen were transfected using lipofectamine according to the manufacturer’s instructions. Green fluorescence signal was observed by fluorescence microscopy at 48h after transfection, and Fluorescence intensity was also tested according to the method described above.

### Statistical Analysis

The experimental data were analyzed using SPSS v19.0 and GraphPad Prism 9 software. The single variance analysis (ANOVA) were used to analyze the differences between groups. Duncan method was used for comparison and P < 0.05 indicated statistical significance. All data are from at least three independent experiments,and expressed as mean ±SEM.

## Results

### Design strategy for auto-circularizing RNA

To assess whether different human tRNA sequences affect circularization of acRNA, we designed a dual reporter gene system that includes green fluorescence and luciferase. The expression of circular RNA depends on the ability of IRES to be recognized without 5’ cap and 3’ polyA tail. Therefore, we constructed the common CVB3-IRES between the 5′ ribozyme and the reporting system. The circularization of acRNA was preliminarily determined by detecting fluorescence and fluorescein intensity. As shown in Fig. 1A, we designed different 5′ ribozymes and 3′ ribozymes based on the sequences from human tRNAs with introns (V1∼V6). Based on the same tRNA, the 5′ ribozyme was placed between the T7 promoter and CVB3-IRES, and the 3′ ribozyme was placed between the reporting system and the linearized site. To confirm the function of the self-shearing design, the cleaved small fragments (from 5’ and 3’-ends respectively) were examined using capillary electrophoresis. From Fig.1B, two corresponding small fragments were detected successfully.

**Figure 1.**
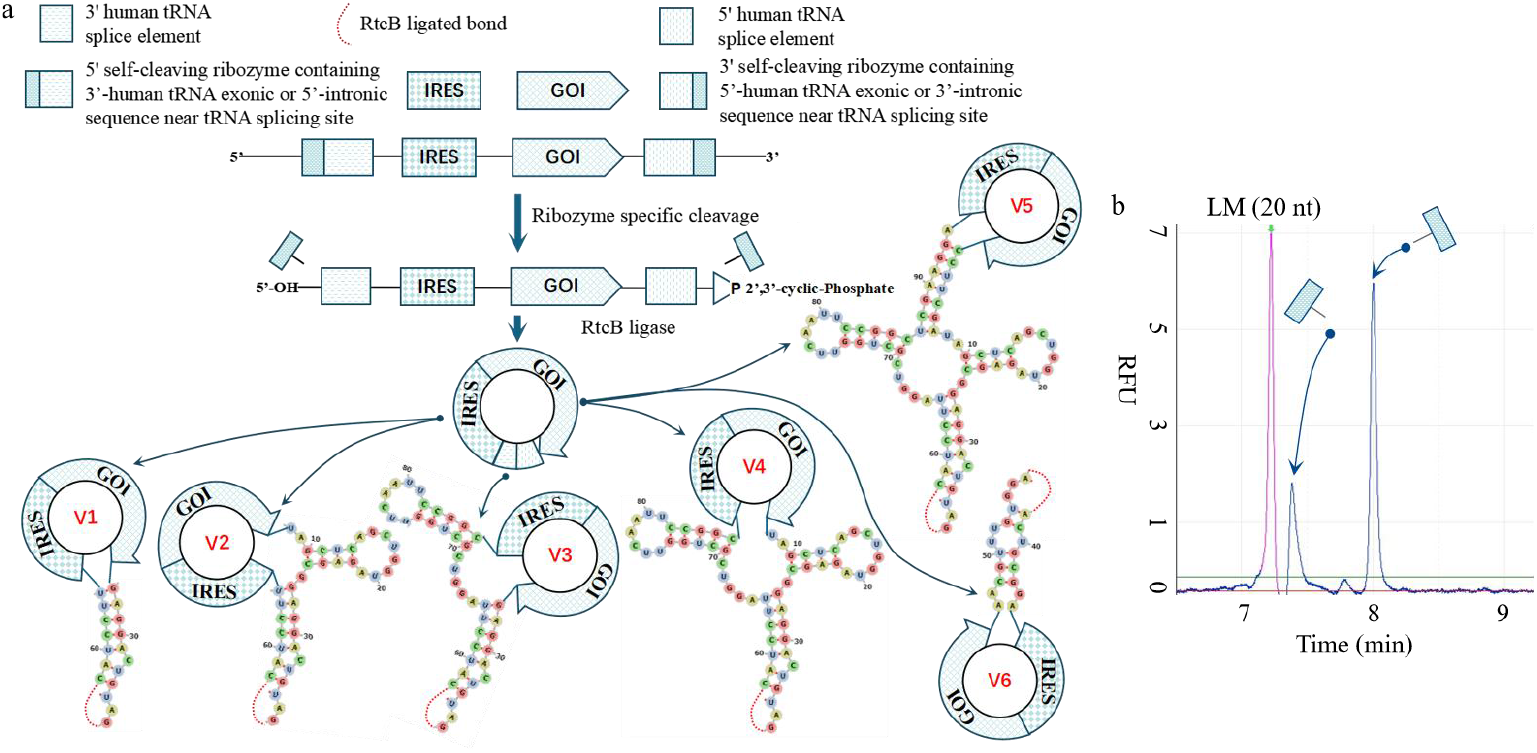
Design acRNA for protein expression based on human tRNAs with introns. (a) The schematic diagram of acRNA designed based on human tRNAs with introns. According to the structure of human tRNAs, six versions (V1-V6) were designed. We ligated the same IRES as well as GOI to different self-splicing elements to form acRNA by end-to-end ligation of intracellular RtcB. The levels of cyclization in different versions were evaluated by reporter genes. (b) Self-cleaved small fragments were detected by capillary electrophoresis. Using 20 nt low marker as a benchmark, two small fragments were detected by capillary electrophoresis at a low voltage of 2 kv.

Based on these results, HEK293T cells were transfected with acRNAs based on the different splicing sequences from human tRNAs with introns (V1∼V6). As shown in Fig. 2A, after 24 h, the significant the green fluorescence signals (NeonGreen) were observed in acRNA groups especially the IleTAT group. This meant that this type of acRNA can be successfully circularized inside the cell. Meanwhile, different tRNA sequences demonstrated different cellular circularization efficiency. From Fig. 2A and 2B, it could be seen that the acRNA designed based on the complete sequence of IleTAT had better circularization effect in vivo compared with other acRNAs. Based on the above results, we selected TyrGTA and IleTAT and further explored whether the acRNAs could be optimized based on their structures. As shown in Fig. 3A, we split the two tRNAs’ sequences according to their structure to construct multiple versions of acRNA. Six versions of acRNA were thus obtained based on TyrGTA or IleTAT tRNAs. V1-V5 was designed based on tRNA exon sequences and V6 was designed based on tRNA intron sequence. By comparing the fluorescence and bioluminescence intensity (NeonGreen-teLuc), as shown in Fig 3B and 3C, the V1 and V2 acRNA based on TyrGTA exhibited better intracellular circularization efficiency than other TyrGTA isoforms. In contrast, the V4 acRNA based on IleTAT exhibited better intracellular circularization efficiency than other IleTAT isoforms.

**Figure 2.**
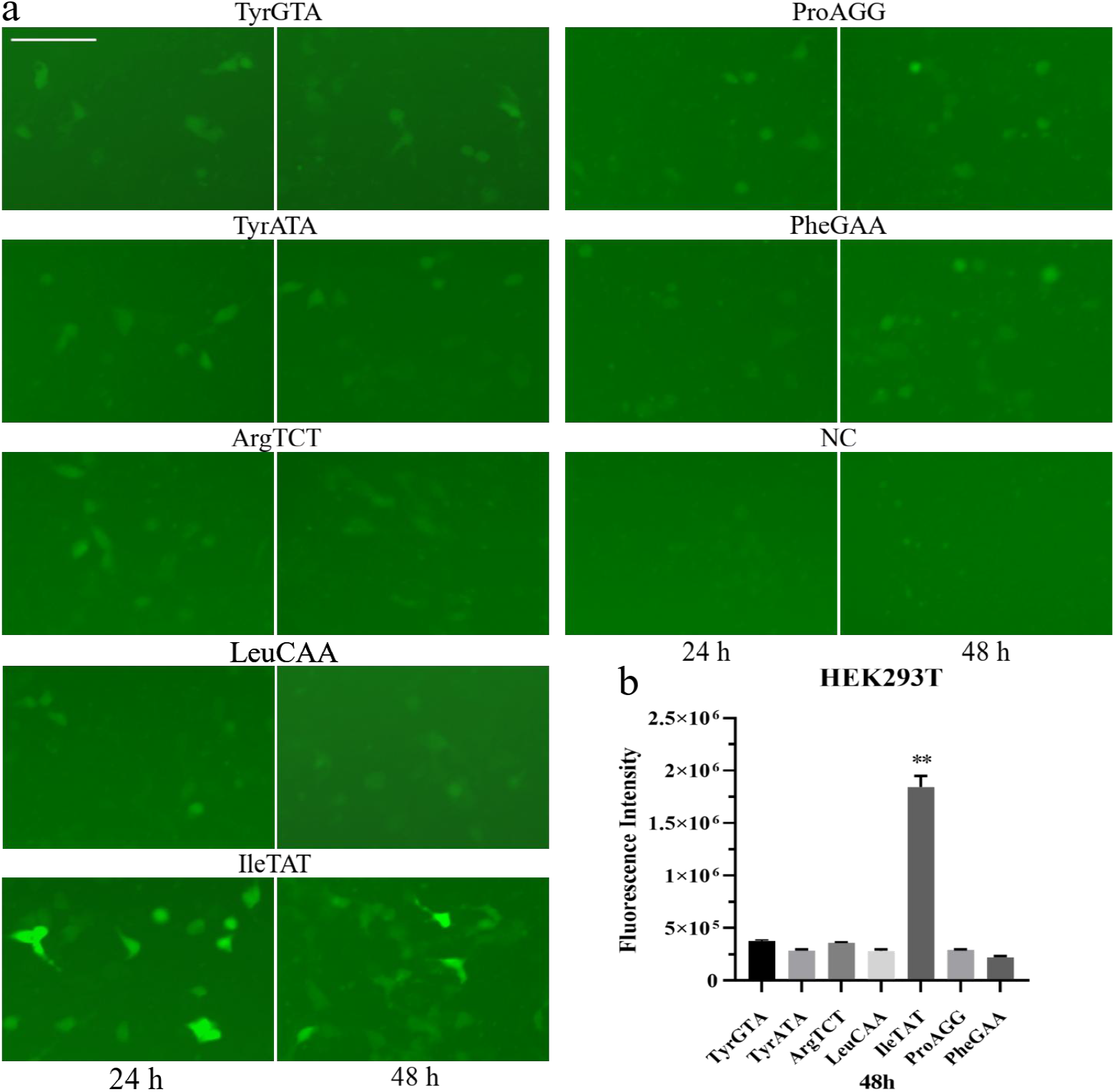
acRNA designed based on the full-length sequences of different human tRNAs in HEK293T cells. (a) The green fluorescence of HEK293T cells transfected with acRNA was observed by fluorescence microscopy at 24h and 48h, respectively. The full-length sequences of TyrGTA, TyrATA, ArgTCT, LeuCAA, IleTAT, ProAGG and PheGAA were selected for in vivo acRNA design for cell transfection assay. RNA without a tRNA-based design ribozyme was used as a negative control (NC). The length of the ruler was 2 μm. (b) The bioluminescence intensity of HEK293T cells was measured at 48 h. The bioluminescence signals of all experimental groups were subtracted by the background bioluminescence from the NC group. Data are presented as means ± SEMs (n=5; ^**^p < 0.01 compared with other groups)

**Figure 3.**
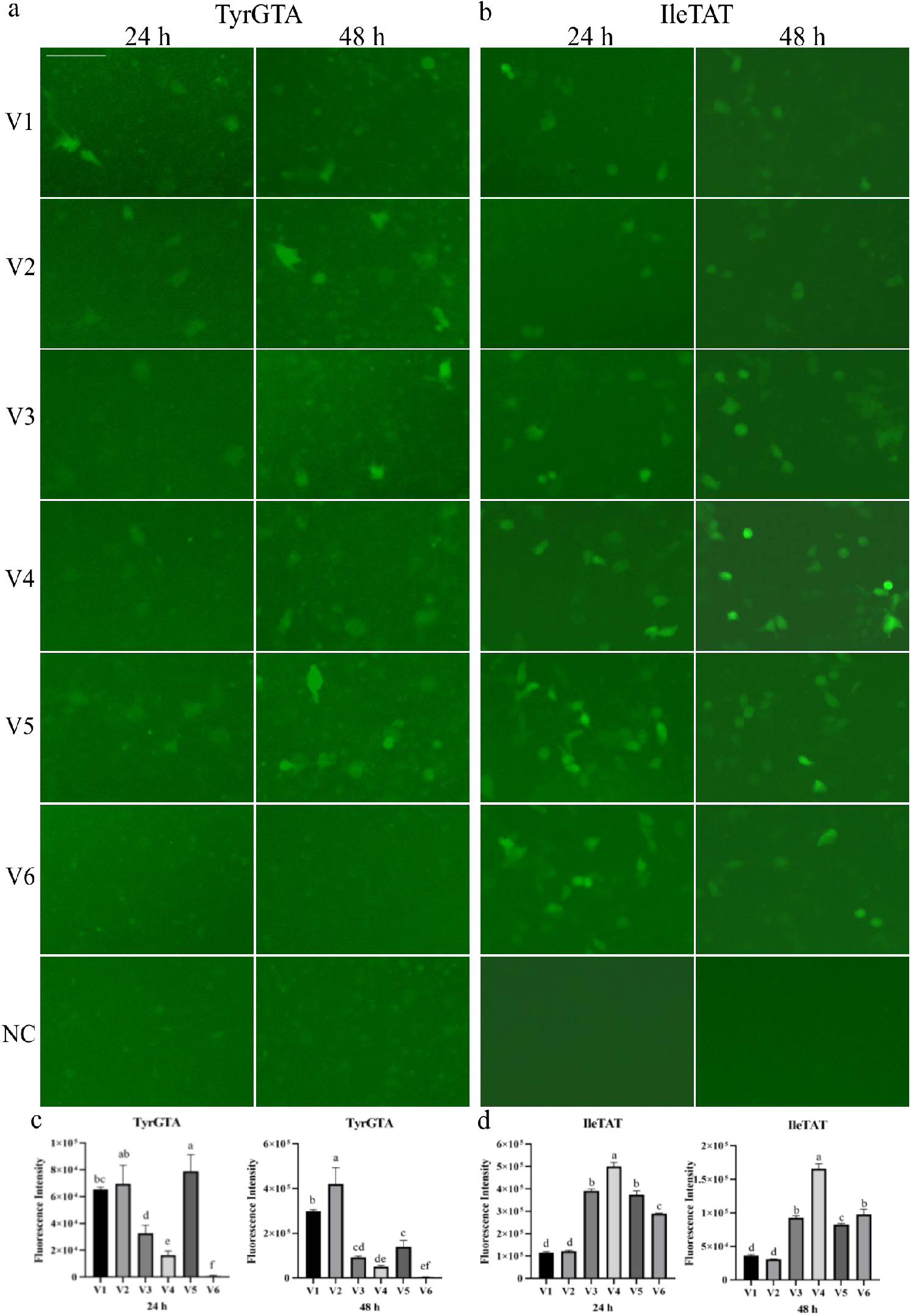
HEK293T cells were transfected with a variety of acRNAs designed based on human TyrGTA or IleTAT sequences. (a) Transfection of acRNA based on the TyrGTA sequence design in HEK293T cells was observed at 24 h and 48 h. RNA without a tRNA-based design ribozyme was used as a negative control (NC). The length of the ruler was 2 μm. (b) Transfection of acRNA based on the TyrGTA sequence design in HEK293T cells was observed at 24 h and 48 h. RNA without a tRNA-based design ribozyme was used as a negative control (NC). (c) The bioluminescence intensity of transfected TyrGTA-based acRNA into HEK293T cells was measured at 24 h and 48 h. d The bioluminescence intensity of transfected IleTAT-based acRNA into HEK293T cells was measured at 24 h and 48 h. The bioluminescence signals of all experimental groups were subtracted by the background bioluminescence from the NC group. Data are presented as means ± SEMs (n=5; Different lowercase letters indicate significant differences at the 0.05 level)

### Further validation of acRNA circularization in vivo

To determine whether acRNA entry into the cell can be circularized normally and whether the ribozyme self-cleavage site is specific, we selected TyrGTA-based V1 and V5 and IleTAT-based V4 for further detection. Each acRNA was transfected into HEK293T cells for 48h, and total RNA was extracted. After specific reverse transcription and PCR amplification, the fragment size was detected using agarose electrophoresis. As shown in Fig. 4A, the predicted bands of several acRNAs circularized in vivo could be detected. We selected the bands amplified by TyrGTA V1 among them for gel recovery and sequencing. The fragment was sequenced from both forward and reverse directions. As shown in Fig. 4B, the marked position of the vertical scissors was the self-cleaving position of the ribozyme. The sequencing results showed that the ribozyme cleaving position was specific and could be successfully circularized in vivo.

**Figure 4.**
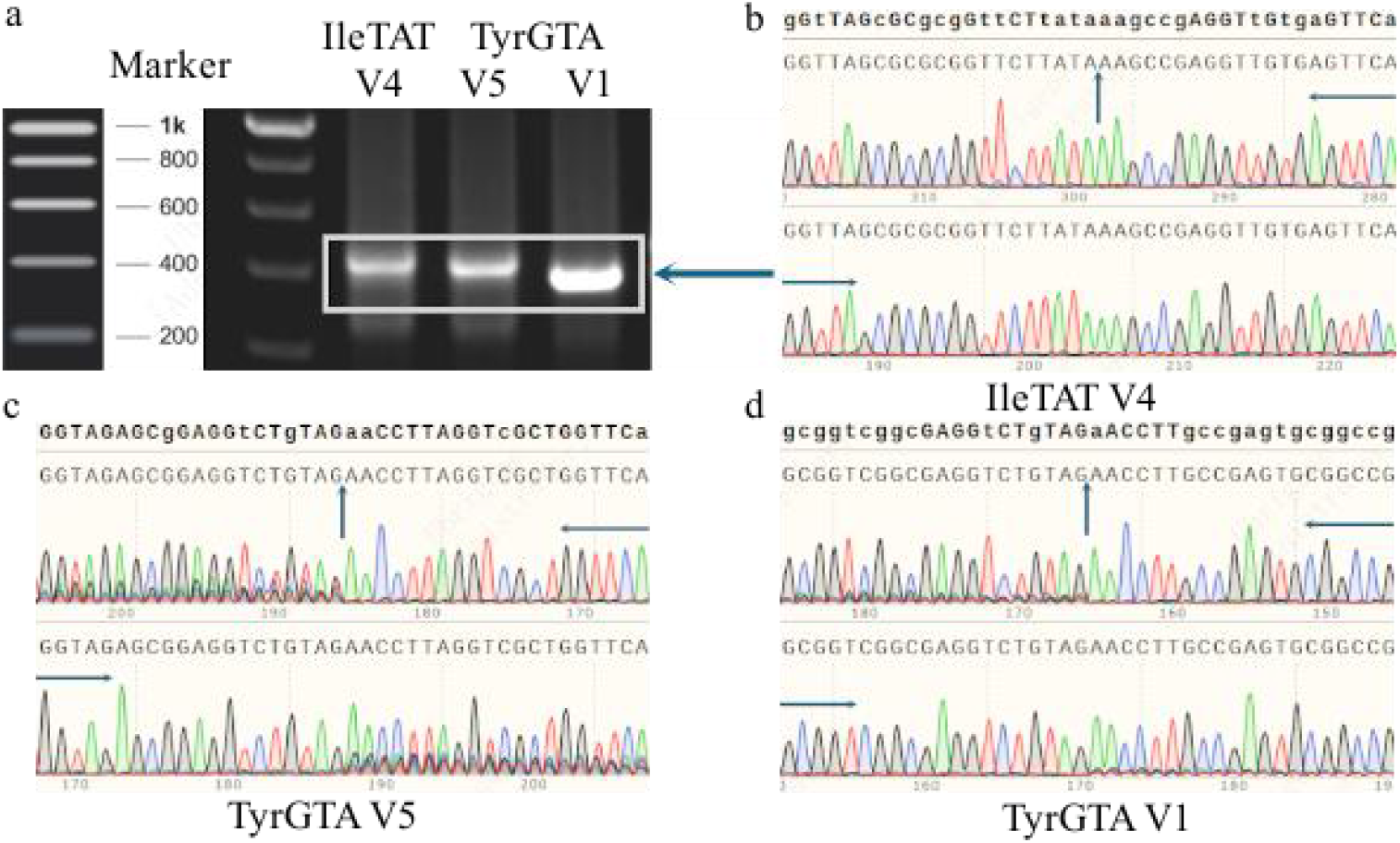
Cyclization verification of acRNA designed based on human tRNA sequences. (a) The looped linker of acRNA was detected using agarose gel electrophoresis. No band could be detected when acRNA was not looped in the cells. The theoretical amplified fragment size of V4 based on IleTAT sequence was 437 bp after in vivo cyclization. At the same time, the theoretical amplified fragments of V5 and V1 based on TyrGTA sequence were 427 bp and 389 bp after in vivo cyclizing, respectively. (b) Sequencing results of the V4 end to end junction based on the IleTAT design. The vertical arrow represents the end to end junction, and the horizontal snip represents the sequencing direction. Other sequencing profiles were the same. (c) Sequencing results of the V5 end to end junction based on the TyrGTA design. (d) Sequencing results of the V1 end to end junction based on the TyrGTA design.

### acRNAs were circularized inside CHO cell and HeLa cell

To verify that acRNA can be circularized in a variety of cells, we selected CHO cells and HeLa cells for acRNA transfection. Based on previous experimental results, we selected the acRNA of TyrGTA V1, V2 and V5. As can be seen from Fig. 5A and 5B, green fluorescence could be observed in CHO cells and HeLa cells after transfection of various acRNAs. Moreover, TyrGTA V1 circularized more efficiently in HEK293T, CHO or HeLa cells in vivo.

**Figure 5.**
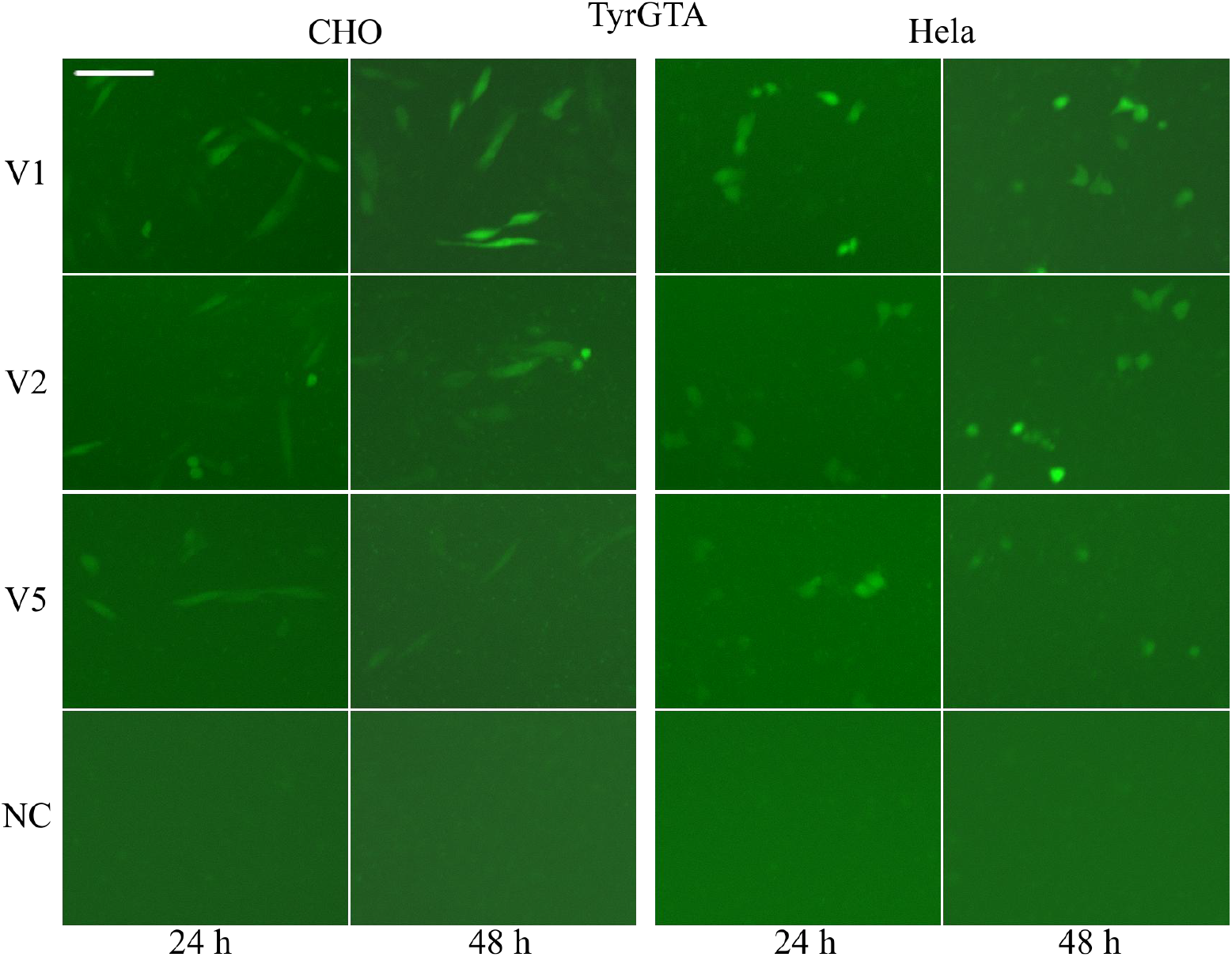
V1, V2 and V5 designed based on the TyrGTA sequence were transfected into CHO cells or HeLa cells. RNA without a tRNA-based design ribozyme was used as a negative control (NC). The green fluorescence of cells was observed at 24h and 48h. The length of the ruler was 2 μm.

### The lower immunogenicity of acRNA

To assess the immunogenicity of acRNA, linear mRNA, acRNAs, and circRNA (circularized *in vitro* by PIE method) were selected for comparison. Total RNAs were extracted from HEK293T transfected for 48 h, and the expression of multiple immune-related genes were detected by reverse transcription and real-time PCR. As shown in Fig. 6, linear mRNA significantly elevated the expression of RIGI, EIF2AK2 and TNF compared with acRNAs and circRNA. acRNA demonstrated comparable or even lower immunogenicity comparing to PIE circRNA, especially for the CCL2 gene. Meanwhile, there was no significant difference in the expression of immune-related genes among the different versions of TyrGTA-based acRNA.

**Figure 6.**
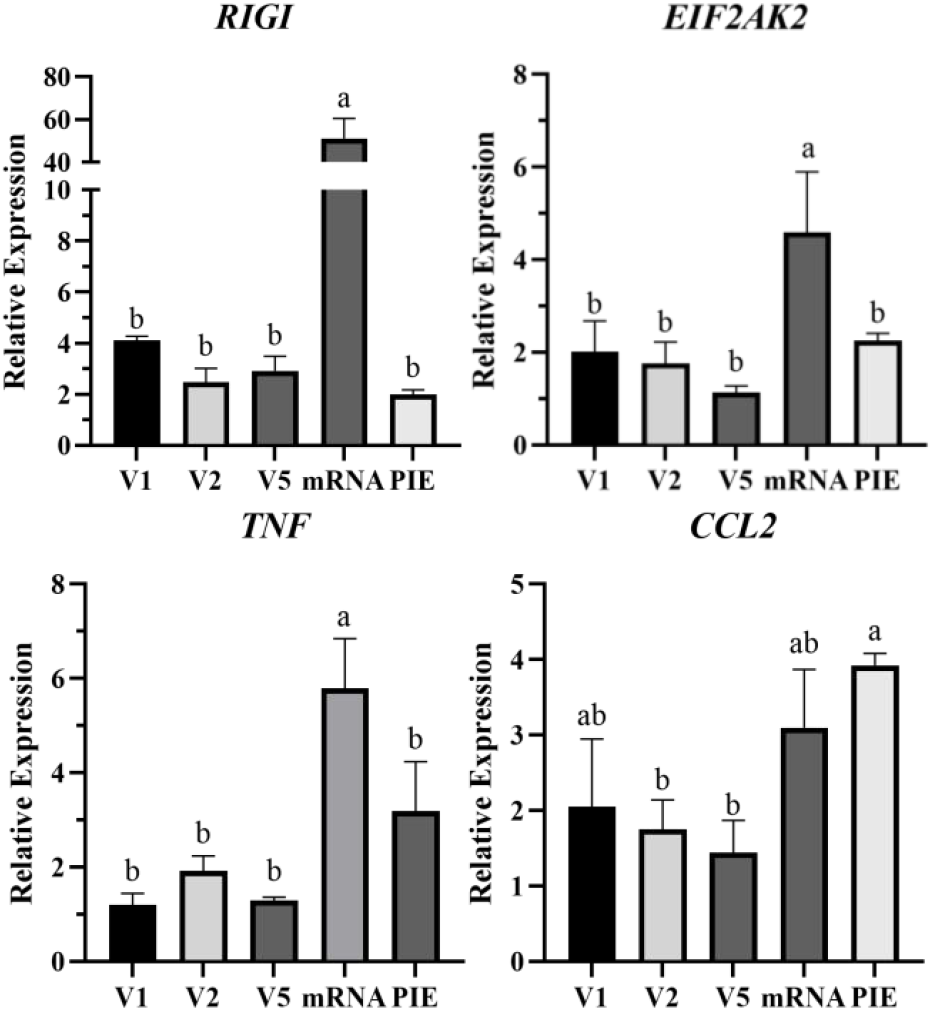
Effect of exogenous RNA on the expression of immune-related genes in HEK293T. We selected several acRNAs designed based on the TyrGTA sequence and compared them with mRNA and PIE looped RNA. RIGI, EIF2AK2, TNF and CCL2 genes were used to assess the intracellular immune response of various RNAs. GAPDH was used as an internal control. Each treatment was performed in triplicate, with two parallel replicates per replicate. Data are presented as means ± SEMs. Different lowercase letters indicate significant differences at the 0.05 level.

### Gene editing with acRNA

Given that our acRNA can be circularized in cells, we sought to explore whether longer sequences could be circularized and expressed in vivo. Cas9-acRNA with a length of ∼5 kb was constructed and transfected into 293T cells with green fluorescence by LNP to explore its gene editing activity. If gene editing is successful, green fluorescent protein expression is disrupted, and a specific CD30 sequence is inserted. As shown in Fig. 7A, the negative control (NC) 293T cells could observe strong green fluorescence, and the addition of Cas9-acRNA or Cas9-mRNA could perform gene editing, that is, reduce the number of cells with green fluorescence. We also measured the fluorescence intensity at the same time and obtained consistent results (Fig. 7B). Although the editing effect of linear mRNA was stronger than that of acRNA, long acRNA was able to expressed large protein (e.g. Cas9) in cells.

**Figure 7.**
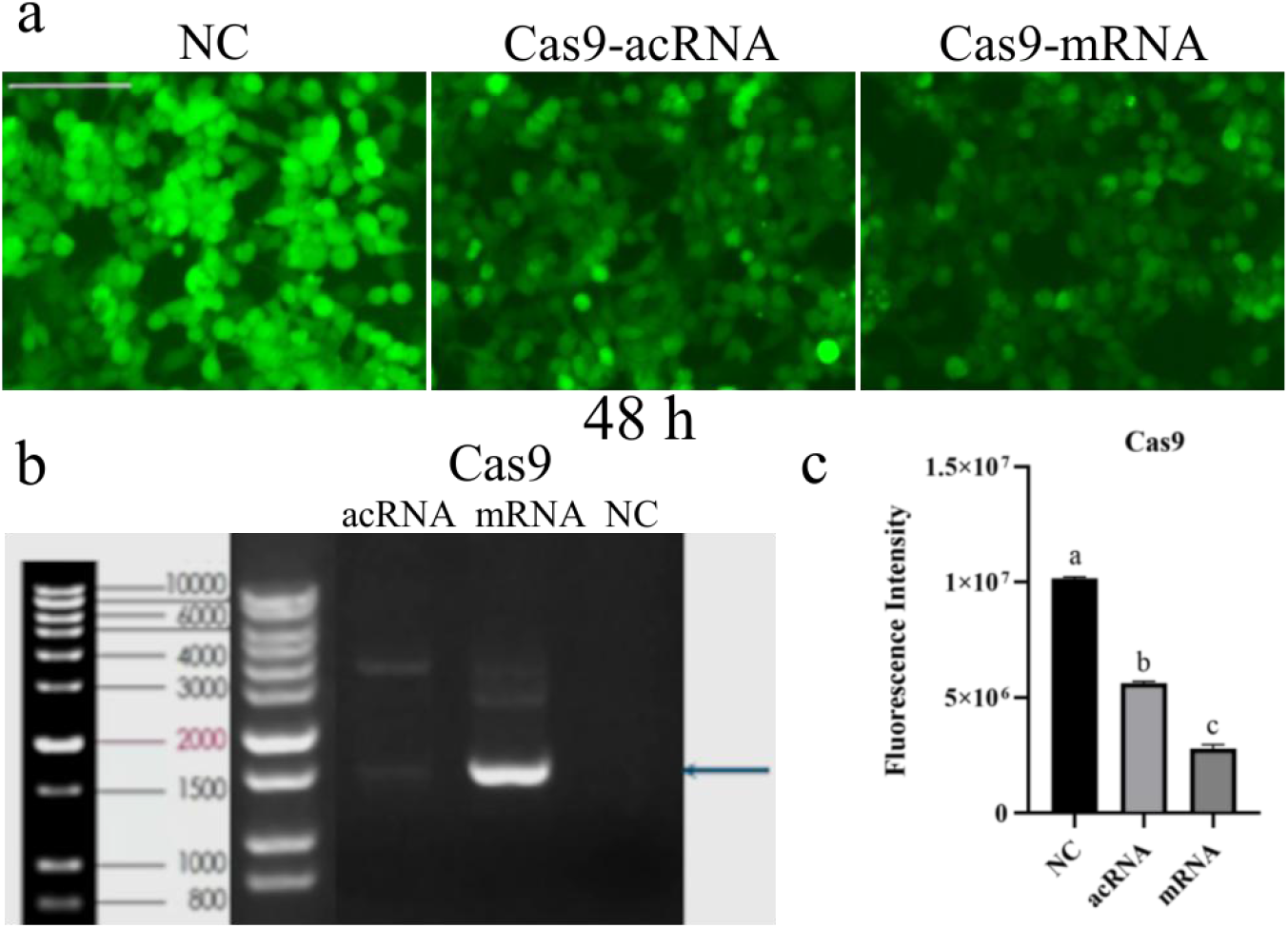
Gene editing was performed using acRNA. (a) The gene editing effects of different RNAs were evaluated by cell fluorescence / bioluminescence reporters. The NeonGreen-teLuc stable expressing 293 cell line were used as a negative control (NC). The length of the ruler was 2 μm. (b) Fragments inserted by gene editing were examined by agarose gel electrophoresis. The marked position of the snip was the location of the band if successful insertion happened. (c) The bioluminescence intensity of cells was detected at 48 h after gene editing. Decrease of the bioluminescence signal meant the NeonGreen-teLuc expressing cassette was destroyed by editing events. Data are presented as means ± SEMs (n=5; Different lowercase letters indicate significant differences at the 0.05 level).

### The longer sustained expression of acRNA in Jurkat cells

To assess the sustained intracellular expression of acRNA, we selected an acRNA (TyrGTA V1) and compared it with an mRNA capable of expressing the same protein. acRNA and mRNA were transferred into Jurkat cells by LNP respectively, and compared with the empty LNP (negative control). The luciferase activity in the cells was then measured for up to ten days. As shown in Fig. 8, although the bioluminescence intensity of mRNA expression in the cells was significantly higher than that of acRNA within the first 48 h, acRNA sustained longer expression of the reporter in Jurkat cells. Moreover, from 72 h to 168 h, the expression of acRNA was significantly higher than that of mRNA in Jurkat cells.

**Figure 8.**
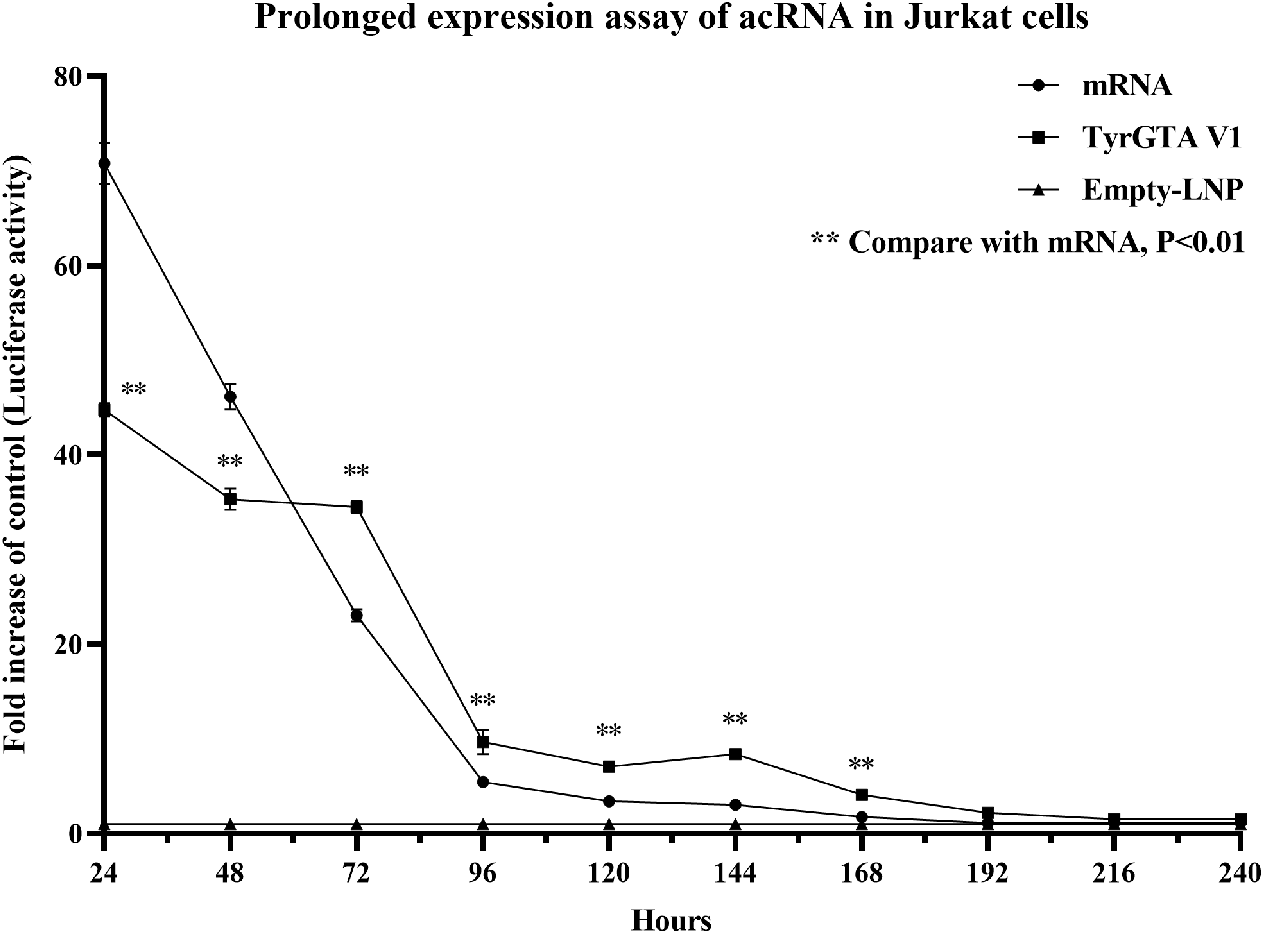
acRNA expresses protein for a longer time than mRNA in Jurkat by LNP. The acRNA and mRNA expressing NeonGreen-teLuc were delivered into Jurkat cells by LNP to compare the duration of their expression and the empty LNP was used as a negative control. The bioluminescence intensity of cells were measured for up to 10 days. Data are presented as means ± SEMs (n=5; ^**^ represents significant difference between acRNA and mRNA, and p<0.01).

## Discussion

CircRNAs have garnered significant attention due to their regulatory functions in gene expression and development^[2, 20]^. The formation of circRNAs involves various mechanisms, including exon back-splicing, group I intron self-splicing, and tRNA intron splicing^[12, 21]^. It has been demonstrated that Metazoans can generate substantial amounts of circRNAs through tRNA intron splicing, which is facilitated by the synergistic action of the tRNA splicing endonuclease (TSEN) complex and RNA ligase^[22]^. The splicing process has been utilized for the development of an *in vivo* expression system to produce tRNA intronic circular RNAs (tricRNAs)^[23]^. However, this system is entirely reliant on endogenous enzymes and exhibits low circularization efficiency. To enhance the system, Jacob et al. utilized the small ribozymes with self-cleavage activity to generate 2′,3′-cyclic phosphates and 5′-hydroxyl termini, which were then connected to form a circular structure *via* endogenous RNA ligase^[19]^. Although this novel approach has been demonstrated to improve RNA aptamer expression levels, the application in protein expression remains unexplored. Furthermore, given that Metazoan tRNA sequences were employed in this system, the safety of RNA vaccines still need to be considered.

In this study, we have successfully developed an efficient and universal intracellular auto-circularization for circRNAs. We have investigated the effect of human tRNA on circRNAs formation and successfully identified effective sequences and tRNA types that enable intracellular auto-circularization. Additionally, we have explored the significance of various factors, including the length of human tRNA splicing elements, ribozyme selection, IRES type and coding gene length, on circRNAs formation. Based on these findings, we established a universal framework template for acRNA and proposed a more efficient and precise method for intracellular auto-circularization. Additionally, we have identified a number of entirely human-derived tRNA splicing sites that can be used in combination with different ribozymes, rendering our approach more flexible than conventional techniques in circRNA synthesis. Research has demonstrated that IRES elements can facilitate the translation of circRNA, and modulate protein synthesis rate^[24-26]^. By utilizing specific IRES elements, we have successfully induced the expression of acRNA and corresponding proteins in targeted cells, thereby enabling tissue or cell-specific gene therapy approaches and delivery systems^[26, 27]^.

Furthermore, a significant breakthrough in our methodology is to synthesize extensive sequences of circRNAs, thereby facilitating stable and efficient expression of larger proteins or functional RNA molecules. Previously, *in vitro* circRNA synthesis was often constrained by the length of coding sequences^[28]^. Although the improved PIE strategy partially addresses this issue, circularization efficiency decreases significantly with increasing fragment length^[16]^. However, our approach demonstrates that it is not constrained by fragment length and can expand the application of circRNA, particularly in the fields of protein engineering and synthetic biology. Furthermore, we have shown that our acRNA constructs maintain their auto-circularization ability via transfection reagents or lipid nanoparticle (LNP) encapsulation. This discovery highlights the resilience and adaptability of our methodology, enabling efficient intracellular auto-circularization regardless of the delivery modality or cell types.

In conclusion, this study presents a novel method for circRNAs generation and establishes an improved intracellular auto-circularization approach. These findings have significant implications for the synthesis and development of efficient and precise gene expression regulatory methods involving circRNAs. Further research and optimization of this technique will contribute to the advancement of RNA technology and potential applications in various fields, including novel RNA vaccines, gene therapy, mimicking endogenous circRNA, gene editing and RNA editing.

