## supplemental table 1 for "Intracellular auto-circularizing RNA using human tRNA splicing elements"

**Supplemental table 1. The tRNA sequences and related ribozymes used in this article**

| Name | Sequence | Description |
|---|---|---|
| Homo_sapiens_tRNA-Tyr-ATA-1-1 | 5’-CCTTCAATAGTTCAGCTGGTAGAGCAGAGGACTATAGctacttcctcagtaggagacGTCCTTAGGTTGCTGGTTCGATTCCAGCTTGAAGGA-3’ | The sequence data of tRNA containing introns are derived from the Genomic tRNA Database. Capital letters represent exons, and lowercase letters represent introns. |
| Homo_sapiens_tRNA-Pro-AGG-3-1 | 5’-GGCTCGTTGGTCTAGGGGTGTGGTTCTCGCTTAGGGaccacagggacaagccCGGGAGACCCAAGAGGTCCCGGGTTCAAATCCCGGACGAGCCC-3’ | |
| Homo_sapiens_tRNA-Arg-TCT-3-1 | 5’-GGCTCTGTGGCGCAATGGATAGCGCATTGGACTTCTAgctgagcctagtgtggtcATTCAAAGGTTGTGGGTTCGAGTCCCACCAGAGTCG-3’ | |
| Homo_sapiens_tRNA-Leu-CAA-2-1 | 5’-GTCAGGATGGCCGAGTGGTCTAAGgcgccagactcaagcttactgcttcctgtgttcgggTCTTCTGGTCTCCGTATGGAGGCGTGGGTTCGAATCCCACTTCTGACA-3’ | |
| Homo_sapiens_tRNA-Ile-TAT-3-1 | 5’-GCTCCAGTGGCGCAATCGGTTAGCGCGCGGTACTTATAagacagtgcacctgtagcaATGCCGAGGTTGTGAGTTCAAGCCTCACCTGGAGCA-3’ | |
| Homo_sapiens_tRNA-Tyr-GTA-7-1 | 5’-CCTTCGATAGCTCAGCTGGTAGAGCGGAGGACTGTAGactgcggaaacgtttgtggacATCCTTAGGTCGCTGGTTCAATTCCGGCTCGAAGGA-3’ | |
| 5' ribozymes based on the human tRNA$^{Tyr}$(GTA) (V1) | 5'-GGGCCGCACTCGCCGGTCCCAAGCCCGGATAAAAAGGGAGGGGGCGGGAAACCGCCTAACCTTGCCGAGTGCGGCCGC-3' | The main sequence design of the ribozymes used in this study. |
| 3' ribozymes based on the human tRNA$^{Tyr}$(GTA) (V1) | 5'-GTGGCCGCGGTCGGC*CCTTCGATAGCTCAGCTGGTAGAGCGGAGGTCTGTAG*AACACTGCCAATGCCGGTCCCAAGCCCGGATAAAAGTGGAGGGTACAGACCTCCG-3' | |
| 5' ribozymes based on the human tRNA$^{Tyr}$(GTA) (V5) | 5'-GGGCCGCACTCGCCGGTCCCAAGCCCGGATAAAAAGGGAGGGGGCGGGAAACCGCCTAACCTTAGGTCGCTGGTTCAATTCCGGCTCGAAGGGCCGAGTGCGGCCGC-3' | |
| 3' ribozymes based on the human tRNA$^{Tyr}$(GTA) (V5) | 5'-GTGGCCGCGGTCGGCCCTTCGATAGCTCAGCTGGTAGAGCGGAGGTCTGTAGAACACTGCCAATGCCGGTCCCAAGCCCGGATAAAAGTGGAGGGTACAGACCTCCG-3' | |
| 5' ribozymes based on the human tRNA$^{Ile}$(TAT) (V4) | 5'-GGGCCGCACTCGCCGGTCCCAAGCCCGGATAAACGGCGAGGGGGCGGGAAACCGCCTAAGCCGAGGTTGTGAGTTCAAGCCTCACGCCGAGTGCGGCCGC-3' | |
| 3' ribozymes based on the human tRNA$^{Ile}$(TAT) (V4) | 5'-TGGCGCAATCGGTTAGCGCGCGGTTCTTATAAACACTGCCAATGCCGGTCCCAAGCCCGGATAAAAGTGGAGGGATAAGAACCGCG-3' | |
