## supplemental table 2 for "Intracellular auto-circularizing RNA using human tRNA splicing elements"

**Supplemental table 2. All primer sequences used in this study**

| ID | Assay | Description | Sequences (5'-3') |
|---|---|---|---|
| cDNA-specific primer | In vivo RNA circularization detection | Reverse transcription primer | CGGACACCCAAAGTAGTCGG |
| CircRNA detection primer F | | Detect whether RNA is circularized | TGGAGGCAGACCGCCGGCTAC |
| CircRNA detection primer R | | | AGGATTAGCCGCATTCAGGG |
| GAPDH F | Real-time fluorescence quantitative detection of gene expression | Evaluation of the intracellular immune response caused by RNA | TGGCACCGTCAAGGCTGAGAA |
| GAPDH R | | | TGGTGAAGACGCCAGTGGACTC |
| RIGI F | | | GCATGGTGTTCCAGATGCCAGA |
| RIGI R | | | TGCTGCTCGGACATTGCTGAAG |
| EIF2AK2 F | | | GGCACCCAGATTTGACCTTCCT |
| EIF2AK2 R | | | TTACTTCACGCTCCGCCTTCTC |
| CCL2 F | | | CCTTCTGTGCCTGCTGCTCAT |
| CCL2 R | | | CTTTGGGACACTTGCTGCTGGT |
| TNF F | | | TCCAGGCGGTGCTTGTTCCT |
| TNF R | | | TGGGCTACAGGCTTGTCACTCG |
